# Antagonizing *cis*-regulatory elements of a conserved flowering gene mediate developmental robustness

**DOI:** 10.1101/2024.11.03.621740

**Authors:** Amy Lanctot, Anat Hendelman, Peter Udilovich, Gina M. Robitaille, Zachary B. Lippman

**Affiliations:** Howard Hughes Medical Institute, Cold Spring Harbor Laboratory, Cold Spring Harbor, NY, USA 11724; Cold Spring Harbor Laboratory, Cold Spring Harbor, NY, USA 11724

**Author notes:** **Corresponding Author** Zachary B. Lippman, 1 Bungtown Rd, Cold Spring Harbor Laboratory, Cold Spring Harbor, NY, USA 11724.

## Abstract

Developmental transitions require precise temporal and spatial control of gene expression. In plants, such regulation is critical for flower formation, which involves the progressive differentiation of stem cell populations within floral meristems followed by rapid sequential development of floral organs. Across plant taxa, these transitions are orchestrated by the F-box transcriptional co-factor gene *UNUSUAL FLORAL ORGANS* (*UFO*). The conserved and pleiotropic functions of *UFO* offer a useful framework for investigating how evolutionary processes have shaped the intricate *cis-*regulation of key developmental genes. By pinpointing a conserved promoter sequence in an accessible chromatin region of the tomato ortholog of *UFO*, we engineered *in vivo* a series of *cis-*regulatory alleles that caused both loss- and gain-of-function floral defects. These mutant phenotypes were linked to disruptions in predicted transcription factor binding sites for known transcriptional activators and repressors. Allelic combinations revealed dosage-dependent interactions between opposing alleles, influencing the penetrance and expressivity of gain-of-function phenotypes. These phenotypic differences support that robustness in tomato flower development requires precise temporal control of *UFO* expression dosage. Bridging our analysis to *Arabidopsis*, we found that although homologous sequences to the tomato regulatory region are dispersed within the *UFO* promoter, they maintain similar control over floral development. However, phenotypes from disrupting these sequences differ due to the differing expression patterns of *UFO*. Our study underscores the complex *cis-*regulatory control of dynamic developmental genes and demonstrates that critical short stretches of regulatory sequences that recruit both activating and repressing machinery are conserved to maintain developmental robustness.

## Main Text

*Cis-*regulatory control of developmental transitions has long been a topic of interest to geneticists—given the broad consistency of somatic genomes across cell types, temporal and spatial regulation of gene expression is the main mechanism by which the functional differentiation of cells that undergirds development is achieved (1). Due to the complexity of gene expression regulation, parsing apart these *cis-*regulatory control nodes has been challenging. While the genomics era has ushered in a host of new strategies to assess transcription, the extreme context-dependence of *cis-*regulation and the highly interconnected genetic interactions that regulate developmental transitions mean that even extensive global analyses of gene expression cannot bridge the gap to phenotype (2). To determine *cis-*regulatory functionality, a combination of genomic contextual data, including chromatin accessibility, transcription factor binding sites, and *cis-*trans interactions, can allow for more precision in our predictions (3).

Conservation across broad evolutionary distances can indicate that genomic sequence is under purifying selective pressure and cannot be mutated away without impeding function and phenotype (4). This has been a guiding principle of molecular evolutionary approaches, which focus on coding sequence conservation as the ratio of synonymous to nonsynonymous mutations can quantify selective pressures. Conserved non-coding sequences (CNSs) can also indicate functionality, and in animal systems deeply conserved non-coding sequences are essential for organogenesis and body plan organization (5, 6). An influx of high-quality annotated genomes has made identification of CNSs across plants feasible (7). These sequences add another informative layer of genomic contextual knowledge to strategies that aim to predict and study *cis-*regulatory functionality.

Regulatory genes that exhibit pleiotropic activity during development are particularly promising candidates to address the still poorly understood relationships between non-coding sequence function and phenotype, as extreme precision in these genes’ expression patterns is indispensable for development (8). Consequently, querying *cis-*regulatory control of these fate drivers can dissect the extent to which these nodes are buffered and how changes in their expression affect the phenotypes they control (9). The regulatory sequences of such genes can impact both penetrance and expressivity of developmental phenotypes, as fine-tuning of their spatial and temporal expression patterns can lead to phenotypic changes of varying severities (10). Penetrance is the genetic concept that a change in phenotype does not manifest in all individuals carrying a particular mutation, both at the organ and organism scale, and expressivity is the related concept that the degree of a phenotype can differ between individuals (11). Despite historical descriptive work on these concepts, the molecular mechanisms behind incomplete penetrance and variable expressivity are poorly understood.

An essential developmental regulator in plants is *UNUSUAL FLORAL ORGANS* (*UFO*), whose pleiotropic roles in flowering and flower formation require exquisite temporal and spatial control of its expression and multilevel function. *UFO* was first identified in Arabidopsis, where null mutants show increased inflorescence branches due to the conversion of floral meristems into secondary shoots (12). Mutants develop flowers that lack petals and stamens, and different mutant alleles show variation in the expressivity of this phenotype, with some alleles causing complete petal loss in all flowers and others showing reduced petal number and homeotic conversions (12). Overexpression of *UFO* increases petal number and flower size (13). These phenotypes align with *UFO*’s expression in the meristem, as *UFO* is first expressed in the transitional meristem that precedes flower formation, and its expression is then spatially restricted to the inner whorls of the floral meristem, promoting petal and stamen development (14).

*UFO* presents a platform in which to study *cis-*regulatory function across evolution, as while its protein sequence and function is broadly conserved across flowering plants, its expression is not. In the Solanaceae, whose primary models for flowering are petunia and tomato, expression of *UFO* orthologs coincides with the floral transition and drives floral identity, whereas in the Brassicaceae *UFO* is also expressed in vegetative tissue (15). Disruption of *UFO* function from classical coding mutations, which impacts all functions in time, space, and level, consequently has more severe phenotypic consequences in Solanaceae species. For example, unlike Arabidopsis null *ufo* mutants, null mutants of the tomato *UFO* ortholog *ANANTHA* (*AN*) fail to make flowers, and instead indefinitely iterate inflorescence meristems and branches (16). Furthermore, transgenic overexpression of *UFO* in the Solanaceae species tobacco (13) and petunia (15) causes extremely early flowering. In tomato, precocious expression of *AN* results in a rapid transition to reproductive growth and single flower inflorescences with large, leaf-like sepals. These phenotypes occur both in transgenic plants where *AN* is driven under the promoters of genes expressed earlier in meristematic maturation and in null mutants of an upstream repressor, *TERMINATING FLOWER* (*TMF*) (17). These strong opposing phenotypes from loss- and gain-of-function imply that *AN* is under exquisite temporal and spatial control. Consequently, *AN cis-*regulation must have evolved to switch between activating and repressing regimes quickly as floral identity is established.

To dissect *UFO cis-*regulation, we used CRISPR to mutate conserved non-coding sequences (CNSs) in the *UFO* and *AN* promoters in their respective species. Perturbation of CNSs within a region of open chromatin in the tomato *AN* promoter strongly affected flowering. Distinct alleles resulted in loss- and gain-of-function mutant phenotypes, suggesting that this region is a hotspot of opposing *cis-*regulatory activity. Perturbation of CNSs in the Arabidopsis *UFO* promoter also affected flower development, with distinct CNSs again giving rise to both loss- and gain-of-function mutants. Our study showcases that targeting CNSs can generate allelic diversity revealing functionally dense *cis*-regulatory sequences that modulate penetrance and expressivity of phenotypes essential for both organism fitness and targeted developmental engineering.

## Results

### A region of open chromatin and conserved sequence is a “hotspot” of *AN cis-*regulation

To determine *AN cis-*regulatory sequence functionality, we leveraged our gene-centric ortholog-based alignment approach, Conservatory (7), to identify conserved non-coding sequences (CNSs) in the *AN* promoter. Conservatory categorizes CNSs by degree of conservation, i.e. the phylogenetic status of the species where conservation in orthologous *cis-*regulatory sequence can be identified. The majority of CNSs in the *AN* promoter are conserved across Solanaceae species, but four CNSs are also conserved to other dicot plant families (**Figure 1A**). We used CRISPR-Cas9 genome editing to perturb five large stretches of the *AN* promoter that contained CNSs, targeting each region separately (**Figure S1, Figure 1B**). For four of these regions (**Figure S1**), we did not observe any changes in plant growth, inflorescence patterning, or flower development. This lack of phenotype could be due to inadequate allelic diversity, as limitations and stochasticity in CRISPR guide design and function does not allow for the complete loss of CNSs in these regions. Alternatively, these CNSs may act redundantly with other CNSs or non-conserved regions of the *AN* promoter, as additive and synergistic interactions among cis-elements can generate higher-order interactions that impact phenotype (18).

**Figure 1:**
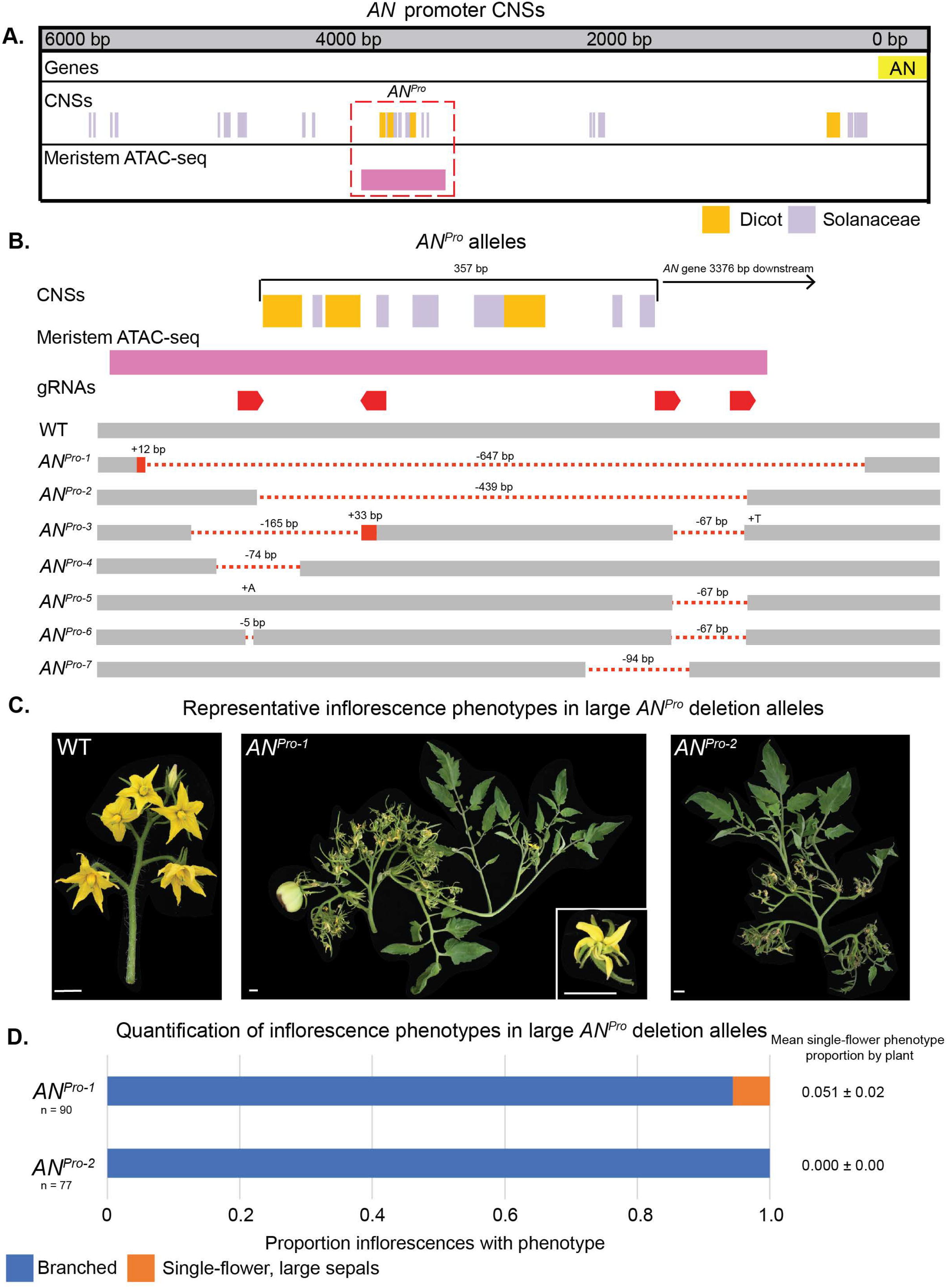
A conserved accessible region of the *AN* promoter is a hotspot for cis-regulatory function. A. Visualization of CNSs within the *AN* promoter sequence. The *AN* promoter is defined as the ∼ 6 kB between the *AN* coding sequence and the proximal upstream gene. CNSs are color-coded by their level of conservation and meristem ATAC-sequencing peaks are depicted. B. Visualization of *AN*^*Pro*^ allelic series. Alleles are ordered by size and location of the deletion. C. Representative inflorescences of WT, *AN*^*Pro-1*^, and *AN*^*Pro-2*^ plants. Scale bars measure 1 cm. D. Quantification of phenotypic frequency in *AN*^*Pro-1*^ and *AN*^*Pro-2*^ plants. Frequency of the single-flower phenotype per plant is given as a mean and standard error.

One promoter region however did show strikingly strong phenotypes when mutated, with different alleles showing distinct effects on development. Notably, this approximately 350 base pair region, three kilobases from the gene body, includes three of the dicot-level CNSs, surrounded by additional Solanaceae-level CNSs (**Figure 1B**). This sequence lies in a region of open chromatin in meristematic tissue as determined by prior ATAC-sequencing, and in fact is the only region of open chromatin in meristems in the entire promoter. CRISPR editing of this region was highly efficient, perhaps due to this chromatin accessibility. The multiple alleles generated allowed us to contrast the phenotypic effects of distinct small sequence perturbations in this CNS-enriched region. We proceeded with in-depth characterization of seven of these alleles, termed the *AN*^*Pro*^ mutants (**Figure 1B**).

### *AN* promoter hotspot mutants affect inflorescence architecture and flower development

Wild type tomato produces multi-flowered inflorescences in a highly stereotyped “zig-zag” pattern of flower initiation (**Figure 1C**), and both genetic and environmental variation can alter this distinctive architecture into more branched forms (18, 19, 20, 21). The largest deletions in the *AN*^*Pro*^ region, *AN*^*Pro-1*^ and *AN*^*Pro-2*^, include a 647 base pair deletion that removes the entirety of the conserved sequence and the majority of the region of open chromatin (*AN*^*Pro-1*^) and a 439 base pair guide-to-guide deletion that removes the entire region of conserved sequence (*AN*^*Pro-2*^). These alleles share a phenotype, proliferatively branching inflorescences that fail to form flowers (**Figure 1C**). These inflorescences do infrequently form flower-like structures that have missing or unfused anther cones (**Figure 1C**), and some develop seed-bearing fruit (**Figure 1C**), although the vast majority do not mature. These phenotypes are similar to a weak coding sequence allele of *AN* (16), indicating that the *AN*^*Pro-1*^ and *AN*^*Pro-2*^ mutants are hypomorphs that show partial loss of *AN* function, likely through the deletion of expression-promoting *cis*-regulatory sequence.

While both *AN*^*Pro-1*^ and *AN*^*Pro-2*^ homozygous mutant plants never form wild type inflorescences, *AN*^*Pro-2*^ plants show complete penetrance of the branched phenotype on all inflorescences, but *AN*^*Pro-1*^ plants do not (**Figure 1D**). Instead, while the majority of *AN*^*Pro-1*^ inflorescences produce iterative branching and malformed flowers, occasional inflorescences on some plants show a striking contrasting phenotype—a single-flower with abnormally large sepals (**Figure 2A**). This phenotype is similar to the *AN* gain-of-function phenotype seen in transgenic plants where *AN* is expressed precociously under gene promoters that are activated in meristem maturation prior to the floral meristem (17). This phenotype is also the hallmark of *tmf* mutants (**Figure 2A)**, which show precocious expression of *AN* in transitional meristems (17). Unlike *tmf* mutants, which exhibit the gain-of-function phenotype on the first-formed inflorescence on the primary shoot, *AN*^*Pro-1*^ plants show this phenotype stochastically throughout plant development.

**Figure 2:**
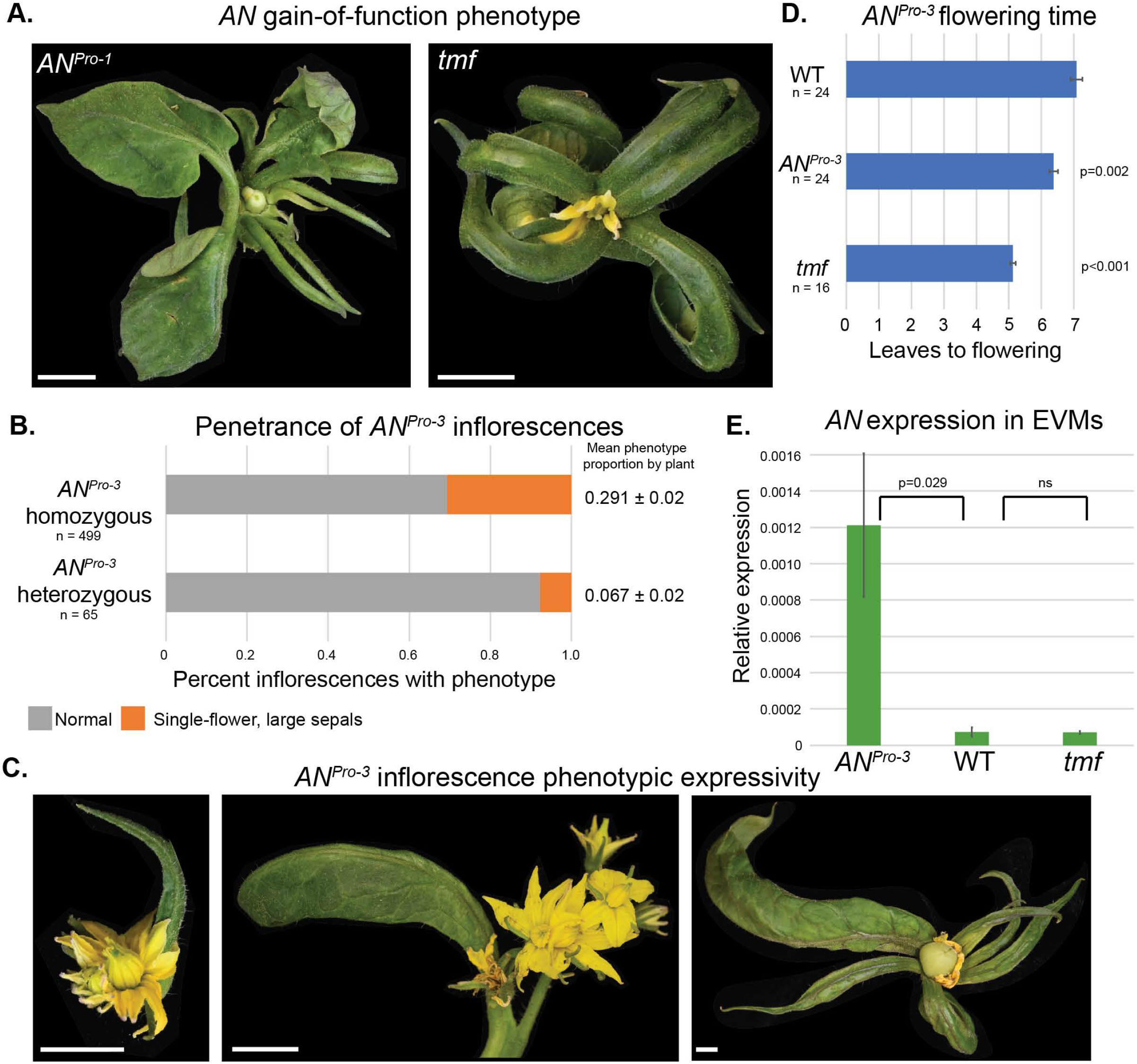
A specific allele of the *AN* promoter hotspot phenocopies gain-of-function *AN* mutants. A. Representative inflorescences of *AN*^*Pro-1*^and *tmf* mutant inflorescences showing the gain-of-function phenotype. Scale bars measure 1 cm. B. Quantification of phenotypic frequency in *AN*^*Pro-3*^ homozygous and heterozygous plants. Frequency of the single-flower phenotype per plant is given as a mean and standard error. C. Representative inflorescences of *AN*^*Pro-3*^ mutants showing the gain-of-function phenotype. Scale bars measure 1 cm. D. Quantification of mean flowering time in WT, *AN*^*Pro-3*^, and *tmf* mutants. Error bars depict standard error. Significant difference from WT was tested via Tukey’s HSD test. E. Quantification of *AN* expression in WT, *AN*^*Pro-3*^, and *tmf* mutant inflorescences at the stage of early vegetative meristems. Significant difference from WT was tested via Tukey’s HSD test.

While *AN*^*Pro-1*^ plants infrequently exhibit this gain-of-function phenotype (**Figure 1D**), it is much more penetrant in *AN*^*Pro-3*^ mutants, with 30% of inflorescences from *AN*^*Pro-3*^ homozygous mutant plants showing this phenotype (**Figure 2B**), while other inflorescences on these plants showed wild type architecture. *AN*^*Pro-3*^ heterozygote plants also show the gain-of-function phenotype, though at a much-reduced frequency, suggesting this allele is dosage sensitive (**Figure 2B**). Phenotypic expressivity is also variable between inflorescences (**Figure 2C**). While all phenotypic inflorescences in both homozygous and heterozygous plants have at least one large leaf-like sepal, some have multiple and the size of sepals vary between inflorescences. Furthermore, these inflorescences are frequently single-flower or have fused flowers. These differences in penetrance and expressivity among plants and between inflorescences within individual plants indicate that *AN* function depends on highly sensitive temporal control to ensure robust inflorescence and floral development, and shows that dosage of this critical developmental regulator can act as a tuning knob in development. This fact that these mutants phenocopy known mutants with aberrant *AN* expression further suggests that these cis-regulatory sequences control repression of *AN*, possibly to prevent precocious *AN* expression. In support, *AN*^*Pro-3*^ plants flower on average of one leaf earlier than wild type plants **(Figure 2D)**, and we found that, unlike WT and *tmf* mutants, *AN* expression is already detectable in early vegetative meristems of *AN*^*Pro-3*^ plants (**Figure 2E**).

### *AN* gain-of-function is caused by deletion of a specific transcription factor binding site

The *AN*^*Pro-3*^ gain-of-function phenotype and early *AN* expression implies loss of repressor activity that allows precocious *AN* expression in early meristematic stages, potentially due to the elimination of repressive transcription factor binding sites (TFBSs). Given that the sequence perturbations in *AN*^*Pro-3*^ are fairly large and complex, we isolated and characterized several smaller deletion alleles. While three alleles (*AN*^*Pro-5*^, *AN*^*Pro-6*^, and *AN*^*Pro-7*^) as homozygous mutants (**Figure 1B**) did not change inflorescence architecture **(Figure 3A)**, *AN*^*Pro-4*^ homozygous mutants (**Figure 1B**) displayed the gain-of-function phenotype (**Figure 3A**). These alleles thus narrowed the region likely responsible for preventing early *AN* expression to a 56 bp sequence surrounding guide one that is completely deleted in *AN*^*Pro-1*^, *AN*^*Pro-3*^, and *AN*^*Pro-4*^ (**Figure 3B**), all of which show *AN* gain-of-function to varying degrees of penetrance and expressivity. While other sequences deleted in these mutants must be important for penetrance (especially as the smallest deletion, *AN*^*Pro-4*^, shows the phenotype the least frequently), this particular region is the only shared deletion among all three alleles. Supporting how loss of this sequence underlies gain-of-function, these 56 base pairs are totally intact in *AN*^*Pro-7*^ and only has one to five base pairs perturbed in *AN*^*Pro-5*^ and *AN*^*Pro-6*^, all of which form wild type inflorescences. Furthermore, *AN*^*Pro-2*^, which is a substantial genome perturbation which never shows the gain-of-function phenotype, also has this region entirely intact (**Figure 3B**).

**Figure 3:**
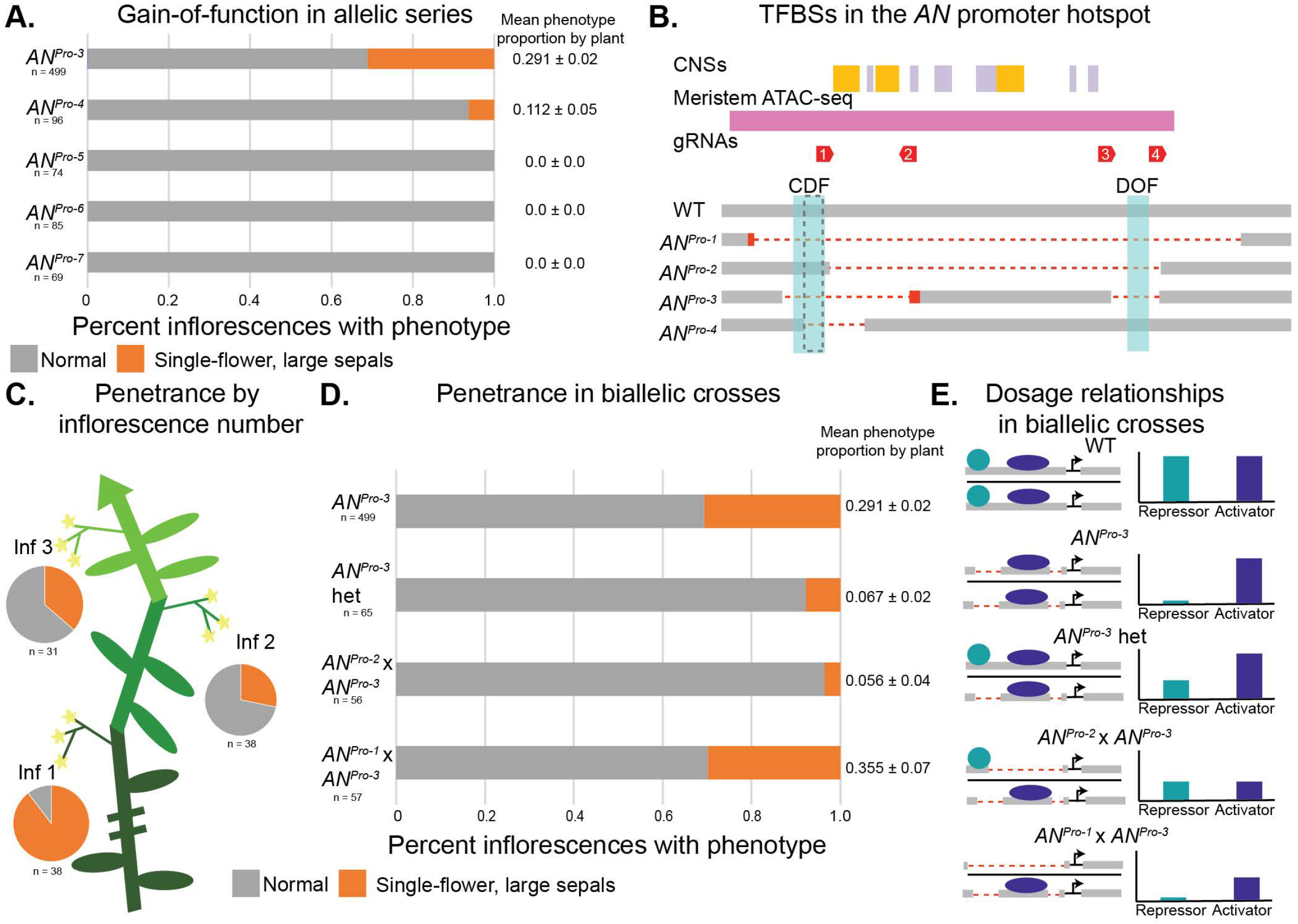
Penetrance and expressivity of the *AN* gain-of-function phenotype depends on genetic lesion, developmental stage and genetic dosage. A. Quantification of gain-of-function phenotype in *AN*^*Pro*^ allelic series inflorescences. Frequency of the single-flower phenotype per plant is given as a mean and standard error. B. Depiction of the *AN*^*Pro*^ hotspot showing putative transcription factor binding sites for DOF transcription factors, highlighted in blue. The 56 base pair region that is deleted in *AN*^*Pro-1*^, *AN*^*Pro-3*^, and *AN*^*Pro-4*^, but intact in *AN*^*Pro-2*^, is outlined by black hatch marks. C. Quantification of phenotypic frequency of *AN*^*Pro-3*^plants by inflorescence number. D. Quantification of phenotypic frequency of *AN*^*Pro*^ biallelic crosses. Frequency of the single-flower phenotype per plant is given as a mean and standard error. E. Proposed mechanism by which biallelic mutants show sensitization or suppression of *AN*^*Pro-3*^’s gain-of-function phenotype by balancing activity of activating and repressing trans-acting factors.

An analysis of TFBSs in the and *AN*^*Pro*^ sequence revealed putative binding sites for distinct transcription factor families. Several overlapping binding sites, immediately proximal to the first guide site itself, are for *CYCLING DOF FACTOR* (*CDF*) family transcription factors (**Figure 3B**). *CDF*s repress precocious flowering in Arabidopsis through suppression of the flowering regulator *CONSTANS*, and Arabidopsis *cdf* mutants flower early in both long and short day conditions (22, 23). There are five *CDF* homologs in tomato, which when expressed in Arabidopsis under a constitutive promoter delay flowering (24), indicating that the ability of *CDF*s to repress flowering is conserved. Our results suggest that in tomato, *CDF*s repress flowering by blocking precocious *AN* expression during meristem maturation, aligning with our finding that *AN*^*Pro-3*^ plants not only show aberrant inflorescence and floral development but also flower early relative to wild type plants (**Figure 2D**). While the *CDF* binding site is partially intact in *AN*^*Pro-4*^, it is completely removed in *AN*^*Pro-3*^, which may explain the difference in phenotypic penetrance between the two alleles. Additionally, there is a putative *DOF* transcription factor binding site deleted in *AN*^*Pro-3*^ that is intact in *AN*^*Pro-4*^ (**Figure 3B**), which may serve as a redundant binding site for CDFs. Indeed, beyond the CDFs, other DOF transcription factors in tomato control inflorescence complexity (25), indicating a potential regulatory role also for DOFs in *AN* function during flower development.

### Penetrance and expressivity of *AN* gain-of-function depends on developmental stage and dosage

The variable penetrance and expressivity of the *AN* gain-of-function phenotype (**Figure 2B, 2C**) shows that there is stochasticity in which inflorescences manifest early *AN* expression or the degree to which this expression impacts phenotype. The penetrance of the gain-of-function phenotype depends on the order in which inflorescences develop on the plant, with phenotypic inflorescences developing earlier and later inflorescences more likely to show normal architecture (**Figure 3C**). Phenotypic expressivity also varies by inflorescence number, with more severe phenotypes, such as single-flower inflorescences with multiple enlarged sepals, emerging more frequently on earlier inflorescences whereas multi-flower inflorescences and flowers with a single enlarged sepal are more frequent on later developing inflorescences. This stochasticity suggests that activity of an *AN* repressor that binds to the deleted sequence can influence penetrance and expressivity, possibly by indirectly influencing the initial maturation states of subsequently formed axillary meristems (26). As tomato is a sympodial system where shoot meristems terminate in floral meristems and new specialized axillary (sympodial) shoots iteratively arise to continue growth, *AN*’s temporal expression patterns in a given transitioning floral meristem can potentially impact the development of the sympodial meristem developing at its base (17). Notably, *tmf* mutant plants also show the gain-of-function phenotype most frequently on the first inflorescence on a plant, with later axillary shoots developing normally (17). These observations are further reinforced by the partial expressivity of the *AN*^*Pro-3*^ allele when heterozygous with an intact functional allele (**Figure 2B**), reflecting a semi-dominant dosage relationship.

To further understand how dosage affects *AN*^*Pro-3*^’s expressivity, we generated biallelic mutant plants between *AN*^*Pro-3*^ and our two hypomorphic loss-of-function alleles: *AN*^*Pro-1*^ and *AN*^*Pro-*2^. We compared the gain-of-function expressivity in these plants to heterozygous *AN*^*Pro-3*^ plants (**Figure 3D**). *AN*^*Pro-2*^ x *AN*^*Pro-3*^ biallelic inflorescences exhibit the gain-of-function phenotype very rarely, at a lower proportion than *AN*^*Pro-3*^ heterozygotes, likely because the combined dosage of a gain- and a loss-of-function allele suppresses this phenotype as balanced dosage is re-established (**Figure 3E**). Interestingly, *AN*^*Pro-1*^ x *AN*^*Pro-3*^ plants show the gain-of-function phenotype at a similar rate as *AN*^*Pro-3*^ homozygotes (**Figure 3D**), even though *AN*^*Pro-1*^ homozygotes primarily show loss-of-function morphology (**Figure 1D**). This may be because *AN*^*Pro-1*^, the largest deletion in this region, likely includes deletions for the binding of both transcriptional activators and repressors (**Figure 3B**). The *AN*^*Pro-1*^ x *AN*^*Pro-3*^ genotype prevents binding for both activators and repressors at one allele (*AN*^*Pro-1*^), but for only repressors at the other (*AN*^*Pro-3*^), causing gain-of-function (**Figure 3E**). These results suggest that balanced dosage of activator and repressor activity in the *AN* promoter is essential for stereotyped inflorescence architecture.

### Shared CNSs between tomato and Arabidopsis are dispersed upstream of *UFO* but maintain function

The pronounced gain- and loss-of-function inflorescence phenotypes in the *AN*^*Pro*^ allelic series suggest that a balance between activator and repressor activity in tightly temporally regulated developmental pathways ensures that core regulators function at precise required timepoints in diverse species. Using Conservatory (7), we found that three dicot-level CNSs share homology with regions of the *Arabidopsis UFO* promoter, spanning 140 million years of evolution. While the sequences are similar, the organization of these CNSs has changed substantially, as CNSs that are more juxtaposed in the *AN* promoter are dispersed throughout the *UFO* promoter (**Figure 4A**). Two of the *AN* CNSs that Conservatory identified as shared among diverged dicot species map to a *UFO* CNS ∼3 kbp upstream of the coding sequence, and the third maps to a CNS ∼1.7 kbp upstream (**Figure 4A**). We mutagenized *UFO* CNSs using CRISPR-Cas9 and generated multiple deletions in three regions, designated *UFO*^*Pro-dis*^, *UFO*^*Pro-mid*^, and *UFO*^*Pro-prox*^. In contrast to *AN*^*Pro*^ mutants, none of the *UFO* mutants showed a loss of floral identity. However, all *UFO*^*Pro*^ mutants that impacted CNSs affected petal development, specifically petal number. Importantly, a deletion in a 1 kbp region having no CNSs between the *UFO*^*Pro-dis*^ and *UFO*^*Pro-mid*^ alleles was indistinguishable from wild type plants **(Supp Fig 2A)**, suggesting that these CNSs are strongly informative of *cis-*regulatory function.

**Figure 4:**
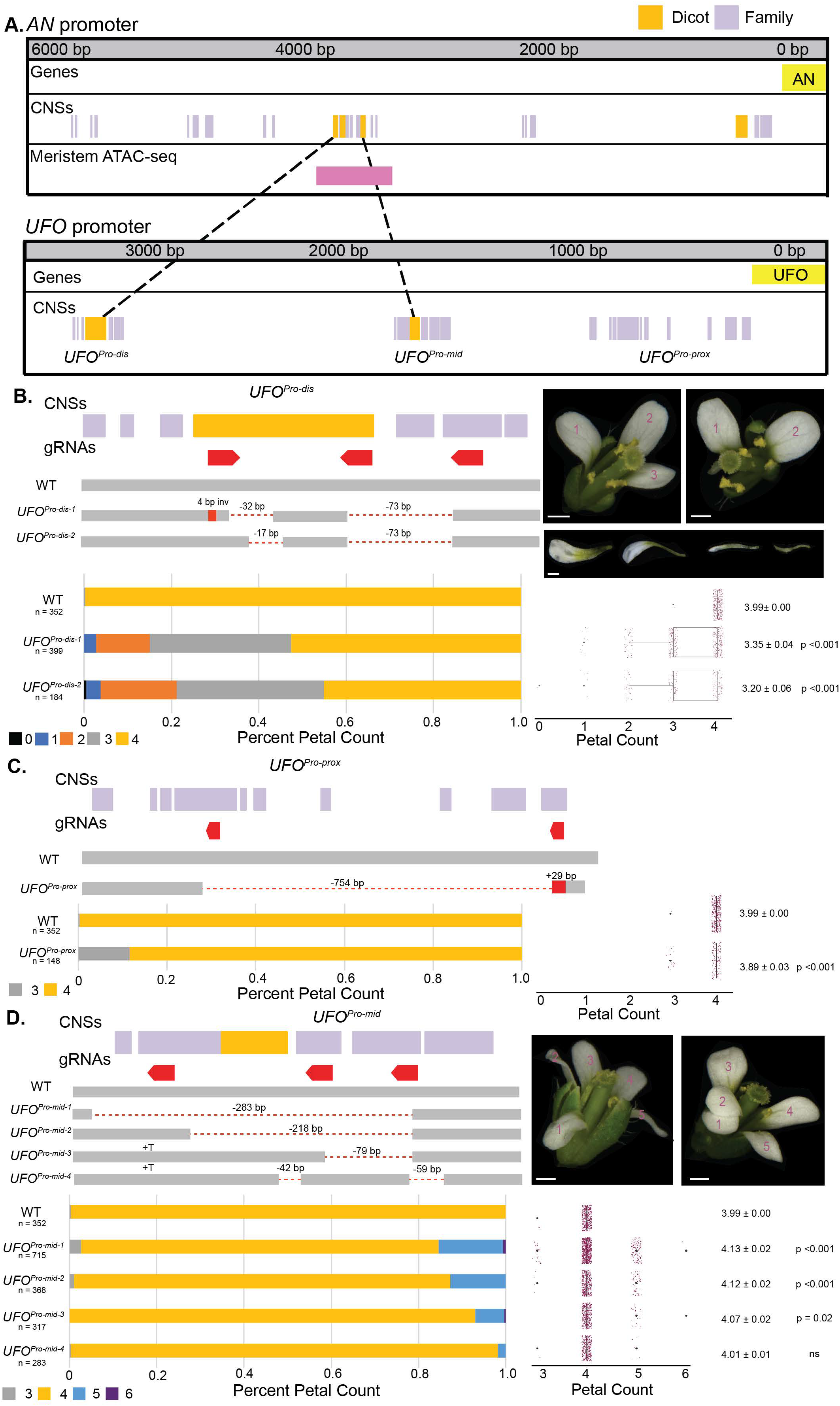
Perturbation of *UFO* CNSs in Arabidopsis breaks petal number canalization. A. Depiction of CNSs in the *AN* and *UFO* promoters and sequence homology of dicot-wide CNSs between the two promoters. B. *UFO*^*Pro-dis*^ alleles are depicted and representative flower and petal images are shown. Scale bars indicate 500 um. Petal counts are depicted as proportions in stacked bars and counts in boxplots. Average petal count and standard error shown. Significant difference from WT was tested via Tukey’s HSD test. C. *UFO*^*Pro-prox*^ allele is depicted. Petal counts are depicted as proportions in stacked bars and counts in boxplots. Average petal count and standard error shown. Significant difference from WT was tested via Tukey’s HSD test. D. *UFO*^*Pro-mid*^ alleles are depicted and representative flower images are shown. Scale bars indicate 500 um. Petal counts are depicted as proportions in stacked bars and counts in boxplots. Average petal count and standard error shown. Significant difference from WT was tested via Tukey’s HSD test.

In Arabidopsis petal number is highly canalized at four petals per flower (27), and decanalization is rare, as most floral homeotic mutants lose petals entirely or aberrantly produce petals in other floral whorls (28). We found that different *UFO* promoter alleles disrupt robustness of petal number in opposite phenotypic directions. In *UFO*^*Pro-dis*^ plants, approximately half of the flowers on a given individual plant form less than four petals (**Figure 4B**), and this difference from wild type was highly significant in both alleles. Strong *UFO* coding mutants lose all petals and stamens, but weaker alleles show variable petal number and petal homeotic conversion (12), suggesting that *UFO*^*Pro-dis*^ mutants are hypomorphs. Interestingly, stamen number is intact in *UFO*^*Pro-dis*^ mutants. This parsing apart of the pleiotropic roles of *UFO* by targeting CNSs may be because specific CNSs control specific developmental processes (7), in this case petal initiation. Alternatively, the distinct processes that *UFO* regulates may be more or less sensitive to quantitative changes in *UFO* expression and function, with the petal whorl being most sensitive to changes to *UFO* cis-regulation during specification of floral meristem and organ identity. Supporting this difference in sensitivity are previously identified insertional mutants in the *UFO* promoter, which also affected petal but not stamen development (29). Interestingly, these mutants lie in a non-conserved region of the promoter downstream of *UFO*^*Pro-dis*^, suggesting it may be the displacement of the *UFO*^*Pro-dis*^ CNS by large T-DNA insertions that drives loss of petals in these mutants.

Petal formation is also aberrant in *UFO*^*Pro-dis*^ plants, with mutants showing often smaller and misshapen petals compared to wild type plants (**Fig. 4B)** This suggests petal initiation is a dose-dependent development process with quantitative outputs—changing the dosage of expression and function of *UFO* can affect petal formation in regards to the shape and size of the petals, not just their number. The variable expressivity among different flowers of *UFO*^*Pro-dis*^ mutant plants mirrors that of *AN*^*Pro*^ mutant inflorescences, and the severity of the petal number decrease in *UFO*^*Pro-dis*^ mutants increases as plants produce more flowers on inflorescence shoots, i.e. flowers that develop later on a given inflorescence have fewer petals (**Sup Fig 2B**). Again, this is similar to the inflorescence age-dependence of the *AN*^*Pro-3*^ mutants’ severity, suggesting that plant age and relative timepoints of inflorescence development can affect the degree to which flower development is able to buffer the impeded function of these CNSs in *UFO* expression.

We observed that *UFO*^*Pro-prox*^ mutants show petal phenotypes similar to *UFO*^*Pro-dis*^ mutants, though much less severe. These plants formed three-petaled flowers much less frequently, though petal number distribution was still significantly different than wild type (**Figure 4C**), suggesting that this CNS also plays a role in promoting petal development. Conversely, *UFO*^*Pro-mid*^ mutants are decanalized in petal number in the opposite direction, increasing petal number to five, and also occasionally forming three-petaled flowers. The more severe *UFO*^*Pro-mid*^ mutants correlated with larger deletions within this CNS, with *UFO*^*Pro-mid-1*^, *UFO*^*Pro-mid-2*^, and *UFO*^*Pro-mid-3*^ all showing a petal distribution significantly different than wild type (**Figure 4D**). These differences between *UFO*^*Pro*^ mutants, much like the opposing phenotypes of *AN*^*Pro*^ mutants, suggest that both activator and repressor transcriptional machinery are binding at these CNSs, leading to precision in the temporal and spatial control of gene expression that promotes robust development. Analysis of TFBSs in the *UFO*^*Pro-mid*^ region identified a putative CDF5 binding site, indicating that there may be shared *cis-*trans regulation of the gain-of-function *UFO* and *AN* phenotypes between Arabidopsis and tomato.

## Discussion

The question of how CNS organization impacts function is of interest to developmental biologists, and a combination of mutational approaches on native sequence (2, 18, 30) and systematic dissection of promoter architecture in synthetic systems (31) are complementary techniques towards a more comprehensive understanding of *cis-*regulatory grammar. Using targeted genome editing of homologous non-coding sequences across broad evolutionary distances, we found that in two highly diverged plant species, conserved sequence is highly predictive of *cis-*regulatory functionality during flowering. While *UFO* CNS sequences share homology between tomato and *Arabidopsis*, their positions relative to the gene body and to each other are different. This so-called “grammar” of CNS positioning clearly affects how these short, conserved sequences exert their regulatory control on their cognate genes (32), as CNSs that are deleted in gain-of-function mutants in tomato (*AN*^*Pro-3*^) instead show a loss-of-function phenotype when perturbed in Arabidopsis (*UFO*^*Pro-dis*^). Recent work comparing Arabidopsis with its close relative *Capsella rubella* found similar shifts in function of conserved cis-regulatory sequence during evolution, due to changes in genomic and developmental contexts (33). It is also undeniable that these allelic series do not encapsulate all of *AN* and *UFO* cis-regulation, as in both species only hypomorphic phenotypes were generated, rather than null mutant phenotypes. With advances in CRISPR-Cas with less stringent requirements for guide targets (34, 35), more precise targeting of CNS sequence can be implemented in the future.

CNSs are likely enriched in TFBSs, as such binding sites are known to be strongly conserved throughout evolution (36). Our TFBS analysis of *AN*^*Pro-3*^ gain-of-function mutants identified binding sites for CDF transcription factors, known repressors of flower formation (24). Analysis of the large region deleted in *AN*^*Pro*^ loss-of-function alleles showed putative binding sites for multiple transcriptional activators, including for MADS-box (37, 38, 39) and MYB transcription factors (40, 41). Intriguingly, a binding site for the AP2 transcription factor AINTEGUEMENTA (ANT) (42) was identified in this region. *ANT* is a known regulator of *LEAFY* (43), *UFO*’s transcription factor co-regulator, so this may be a mechanism by which expression of these two genes are coordinated in tomato, as *ANT* regulation of both *AN* and the *LEAFY* ortholog, *FALSIFLORA* (44), could promote flower formation. Alternatively, given that *AN* is only activated in the floral meristem in tomato, similar to the expression pattern of *LFY* in *Arabidopsis* (15), *AN* and *LFY* may be closer “expression orthologs” than their respective true evolutionary orthologs, due to *ANT* binding to the respective promoters in the two species.

CNS organization is not the only distinction between the *AN* and *UFO* promoters. The phenotypes emerging from the CNS allelic series also differ between tomato and Arabidopsis. *UFO*^*Pro*^ mutants show defects in petal number canalization whereas *AN*^*Pro*^ mutants show defects in inflorescence architecture and flower formation. These different phenotypes reflect divergence in the pleiotropic roles of *UFO* during flower development. In Solanaceae species, *UFO* orthologs drive the transition to flowering (15), causing gain-of-function *AN*^*Pro*^ mutants to promote precocious *AN* expression and to flower early. In contrast, in Arabidopsis, *UFO* is expressed broadly during meristematic development and *LFY* expression instead drives flower formation (15). This may explain why *UFO* CNS gain-of-function mutants do not affect flower formation—as *LFY* expression remains intact in *UFO*^*Pro*^ mutants, flowering is undisrupted. Notably, the sequence that is deleted in all *AN*^*Pro*^ gain-of-function mutants is not in a region of conservation, but rather immediately proximal to a dicot-level CNS. Whether CNS-proximal sequence may be a global repository to encode lineage-specific *cis-*regulatory function (such as *UFO* expression driving flowering in the Solanaceae) remains to be seen. Work in animal systems has shown that *de novo* enhancer formation is more likely to generate phenotypic novelty than changes in conserved enhancer sequences (45), suggesting that non-conserved sequence may more likely to promote developmental divergence across evolutionary time.

In both tomato and Arabidopsis, disrupting CNSs strongly affects overlapping developmental programs that are critical for reproduction, inflorescence architecture and petal number. Petal number in Arabidopsis is incredibly canalized, with near invariant formation of four-petaled flowers across ecotypes and environmental conditions (27). Interestingly, *Cardamine hirsuta*, a close relative of Arabidopsis, has naturally decanalized petal number that is affected both by genetic background (46) and environmental conditions (47). Given that Arabidopsis *UFO*^*Pro*^ mutants can recapitulate Cardamine’s decanalization, it would be interesting to explore how modulating *UFO* expression and function during petal organogenesis can reveal phenotypic variation more broadly across evolution. Though inflorescence architecture and floral organ number are already variable phenotypes in tomato, in part due to domestication (19), ablation of CNSs in the *AN*^*Pro*^ mutants causes even stronger effects on flowering and floral development, reinforcing that CNSs of essential developmental genes are regulatory hubs for canalized development. Arabidopsis and tomato also differ in their inflorescence organization—while Arabidopsis exhibits monopodial growth with inflorescences budding off an indeterminately growing shoot apical meristem, tomato has a sympodial growth habit, with each meristem terminating into a differentiated flower and new growth continuing from specialized (sympodial) axillary meristems (26). These differences in growth habit could contribute to the differences in phenotypic severity between the *UFO*^*Pro*^ and *AN*^*Pro*^ allelic series, suggesting that developmental trajectory differences across evolution affect the phenotypic consequences of modulating *UFO* function.

In both species, CNS-targeting mutants showed incomplete penetrance and variable expressivity both between plants and among inflorescences and flowers within a given individual. This variation suggests that the phenotypic manifestation of these perturbations depends on developmental progression, genetic dosage, and environmental conditions. The incomplete penetrance of *AN*^*Pro-3*^ phenotype in biallelic plants in particular hints to the exquisite dosage-dependence of *AN* expression and function, as the combination of gain- and loss-of function mutations in biallelic mutant plants return to robust inflorescence development, likely due to a rebalancing between activator and repressor activity. These shifts in phenotypic penetrance due to allelic dosage is only visible because the gain-of-function phenotype is semi-dominant and thus present in *AN*^*Pro-3*^ heterozygous plants. These opposing deviations from robustness in distinct alleles shows that regulation of *AN* and *UFO* expression is clearly an inflection point in flower formation across species. CNSs are primed to integrate activator and repressor regimes, the slightest shift between which can cause strong effects on development.

An obvious but challenging next step would be to link the precise molecular consequences of non-coding alleles with their phenotypic penetrance and expressivity. Advances in *in vivo* reporter assays (48) and scRNA-sequencing (49) could elucidate the temporal and spatial expression patterns of transcriptional regulators such as *UFO* in rare cells and developmentally transient tissue types such as maturing floral meristems and connect these expression patterns to the incomplete penetrance and variable expressivity among individual inflorescences and plants. Genomic methods to quantify transcription factor binding (50, 51) could bridge these analyses in *cis*-regulation to the trans-acting factors that bind to these sequences. For all these methods, allelic series, both engineered as in this work (2, 18, 30) as well as those derived from natural variation in the germplasm (52), are prime genetic resources to explore this link between quantitative expression changes in critical developmental regulators and the degree of penetrance and expressivity changes displayed by these *cis*-regulatory alleles (53). The more knowledge gained on these inflection points in other developmental programs and involving other core genes, the more we can understand the underlying molecular inducers of penetrance and expressivity. With current genomic and gene editing tools to mimic natural variation and go beyond it, we can form a more complete picture of how robustness is maintained through the opposing functions of activating and repressing transcriptional machinery (54), and how these *cis-*regulatory regimes can provide unique targets and opportunities for trait engineering.

## Materials and Methods

### Plant material, growth conditions and phenotyping

*Solanum lycopersicum* cv. M82 is the background cultivar for all WT and transformed tomato mutagenesis experiments. Tomato seeds were sown directly in 96-well flats for 4 weeks before being either transplanted to pots and grown in greenhouse conditions or transplanted directly to fields at Uplands Farm at Cold Spring Harbor Laboratory in summer growth seasons. The greenhouse uses natural and supplemental artificial light (from high pressure sodium bulbs ∼250 umol/m^2^) in long-day conditions (16h light, 8h dark) and is maintained at a temperature between 26–28°C (day) and 18–20°C (night), with relative humidity 40–60%. Field-grown plants were grown with drip irrigation and standard fertilizer regimes. For each unique genotype, inflorescence phenotypes were characterized for at least four inflorescences from at least ten plants.

Inflorescence phenotypes were quantified from the first-developing inflorescences on the primary and secondary shoot. For flowering time quantification, plants were grown in greenhouse conditions until flowering. Leaf count before the first inflorescences was quantified for sixteen to twenty-four plants for each genotype. All raw data is described in **Data S1**.

*Arabidopsis thaliana* ecotype Col-0 is the background cultivar for all WT and transformed Arabidopsis mutagenesis experiments. Arabidopsis plants were germinated on 1/2 MS plates and transplanted to 48-well flats for growth. Plants were grown in growth chambers under long day conditions (16h light, 8h dark) at 22°C and light intensity ∼100 umol/m^2^. For each unique genotype, petal number was quantified from at least ten flowers from twelve plants. Petal number was qualified for the first-developing flowers on the primary shoot. All raw data is described in **Data S1**.

### CRISPR-Cas9 mutagenesis, plant transformation, and selection of mutant alleles

Transgenic tomato seedlings were generated via CRISPR-Cas9 mutagenesis as previously described (55). Guides were selected for proximity to CNSs and were designed using Geneious Prime (https://www.geneious.com). Guide RNAs, Cas9, and kanamycin selection genes were cloned into a binary vector via Golden Gate assembly (55, 56). This vector was then transformed into tomato via *Agrobacterium tumefaciens* mediated transformation in tissue culture (55). Transgenic plants were screened for mutations using PCR primers surrounding the gRNA target sites and Sanger sequenced to determine mutant identity. First or second generation transgenic plants were backcrossed to WT and Cas9-negative progeny were selected for phenotypic characterization.

Transgenic Arabidopsis were also generated via CRISPR-Cas9 mutagenesis using the Golden Gate assembly method to clone binary vectors containing the guide RNAs, Cas9, a kanamycin selection cassette, as well as a pFAST-R selection cassette used for seed coat color screening for transformants (56, 58). The Arabidopsis cassette used an intronized Cas9 previously demonstrated to increase editing efficiency (59). Cloning of this cassette is described in (30). Arabidopsis plants were transformed with binary vectors using *Agrobacterium tumefaciens* floral dip (60). Transgenic seeds were selected using fluorescent microscopy and germinated on 1/2 MS plates before transferring to soil at seven days post germination. Initial editing generations (T1 plants from T0 (dipped) parents) were subjected to a heat cycling regime shown to increase Cas9 editing activity (61). Growth chambers were set to shift between 37°C for 30 h and 22°C for 42 h for 10 days, before returning to normal long day conditions. T1 flower DNA was genotyped to identify plants that showed editing. Seeds from these plants were counter-selected by fluorescence for absence of Cas9 and screened in the next generation for mutant identity and zygosity. T3 homozygous plants were phenotyped. All gRNA and genotyping primer sequences are available in **Data S2**.

### *Cis*-regulatory sequence conservation analyses and TFBS prediction

Conserved non-coding sequences were identified via Conservatory (7) and ATAC-sequencing peaks were obtained from previous work on meristem chromatin accessibility in our lab. CNS sequences are listed in **Data S3**. Transcription factor binding sites were predicted within the conserved regions in the *AN* and *UFO* promoters using FIMO in the MEME suite (62). The TFBS position frequency matrices used were acquired from the JASPAR CORE PFMs of plants collection (63). A p-value cutoff of 0.00001 was used to predict TFBSs.

### RNA extraction and quantification of *AN* expression

Seeds of the relevant genotypes were germinated on wet filter paper at 28 °C in the dark, and transplanted to soil in 96-well plastic flats and grown in greenhouse conditions once germinated. Meristems were harvested 5-7 days after transplant after microscopy confirmation of early vegetative meristem (EVM) stage. Thirty meristems per replicate were harvested with three biological replicates per genotype. Meristems were immediately flash-frozen in liquid nitrogen upon harvest and total RNA was extracted using TRIzol Reagent (Invitrogen). Five hundred ng of RNA was used for complementary DNA synthesis with the SuperScript IV VILO Master Mix (Invitrogen). Quantitative PCR was performed with gene-specific primers using the iQ SYBR Green SuperMix (Bio-Rad) reaction system on the CFX96 Real-Time system (Bio-Rad). Primer sequences are available in **Data S2**.

## Acknowledgments

We thank members of the Lippman laboratory for helpful comments and discussions, and assistance with phenotyping; J. Van Eck and Q. Jiang and their respective teams for performing tomato transformations; and T. Mulligan, K. Schlecht, B. Fitzgerald and S. Qiao for assistance with plant care.

## Author Contributions

A.L., A. H., and Z.B.L designed research; A.L, P.U. and G.M.R performed researched; A.L. and Z.B.L wrote the paper.

## Figure Legends

**Figure S1:**
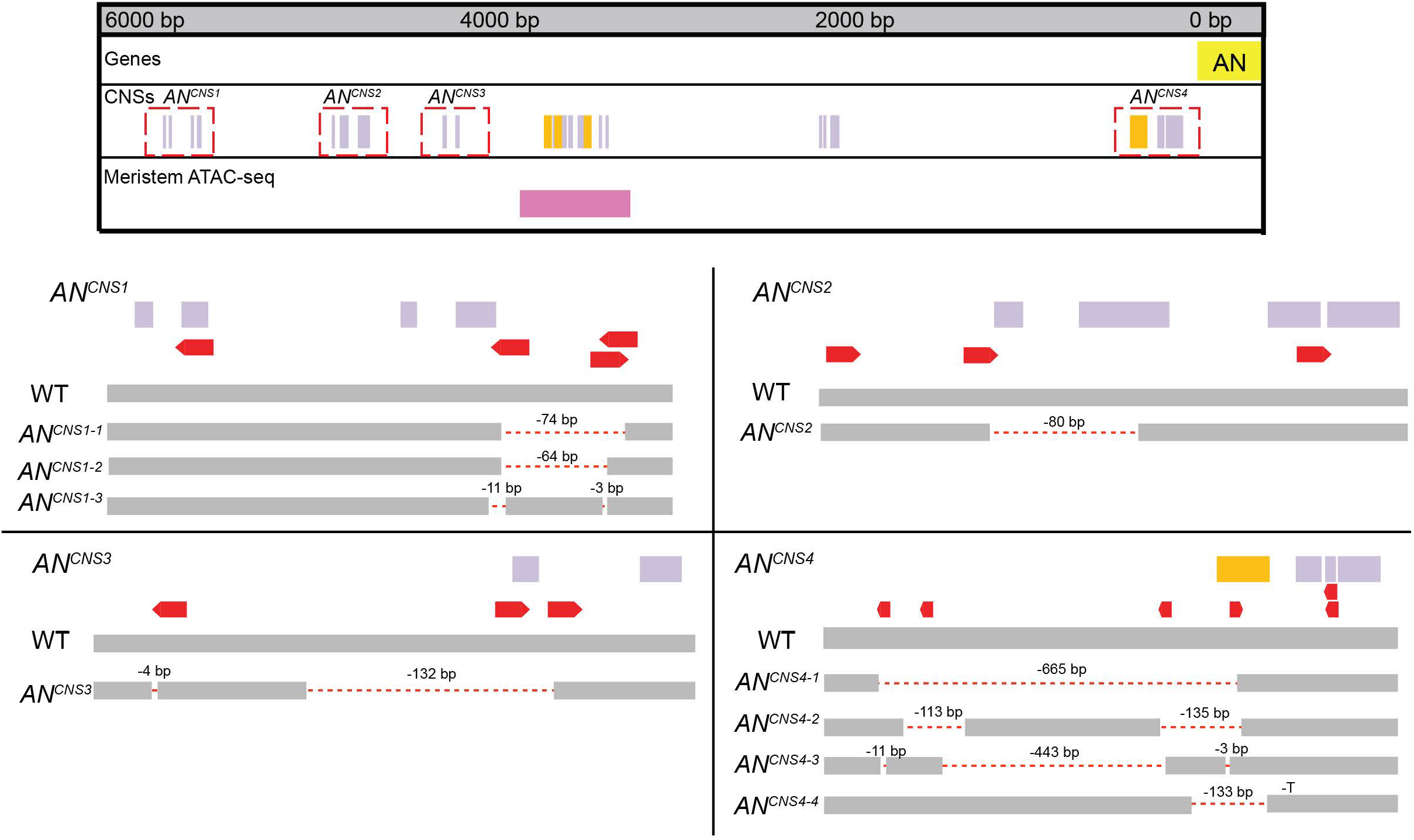
CRISPR targeting of *AN* CNSs. Depiction of CRISPR alleles generated in the four constructs targeting CNSs in the *AN* promoter that are not in regions of open chromatin.

**Figure S2:**
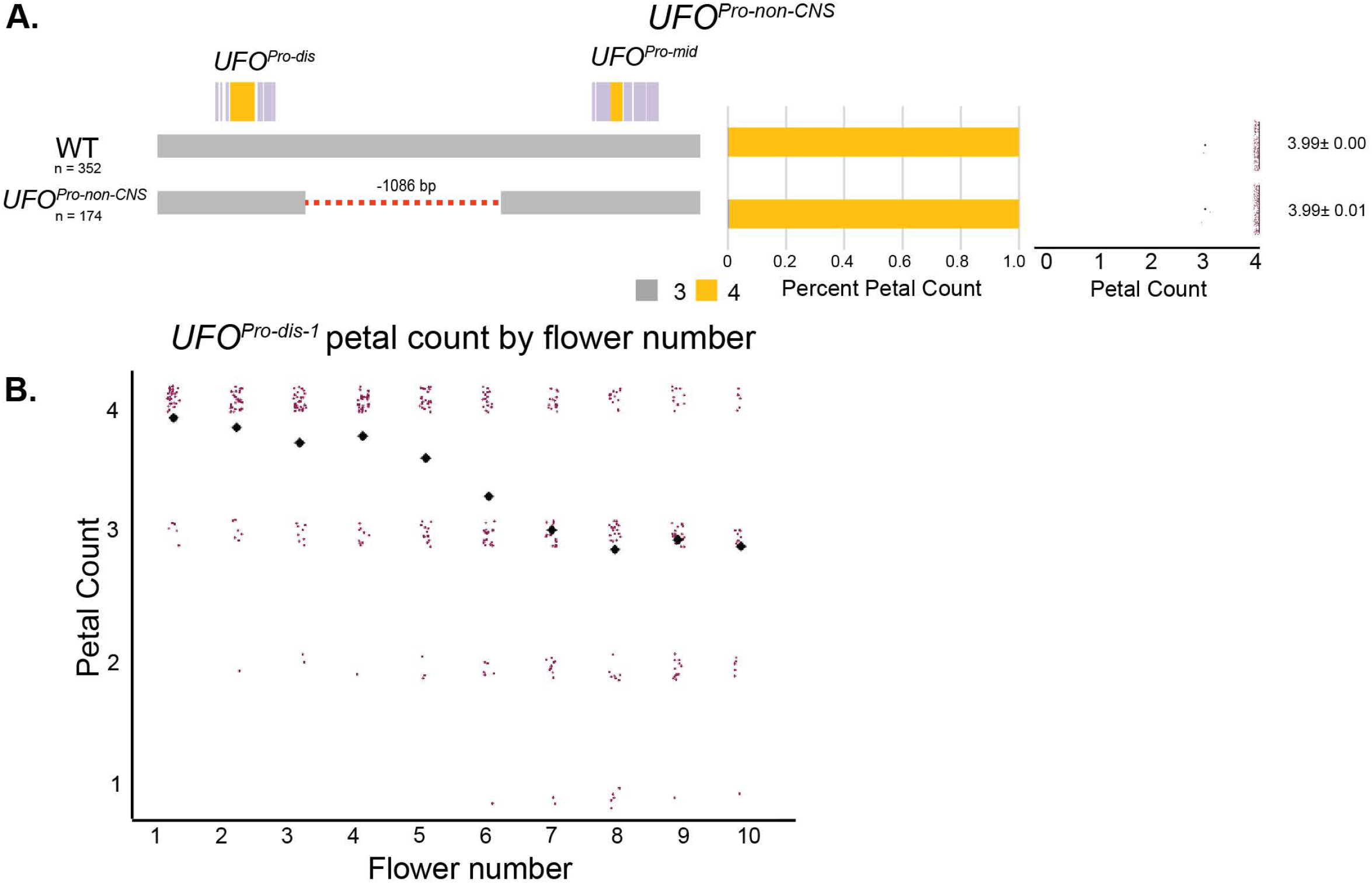
Petal count decanalization depends on genetic lesion and flower number. A. The *UFO*^*Pro-non-CNS*^ allele is a ∼1 kbp deletion between the distal and mid-promoter CNSs. *UFO*^*Pro-non-CNS*^ mutants do not show petal number decanalization or any floral developmental defects. Petal counts are shown as proportions in stacked bars and total counts in boxplots. Average petal count and standard error shown. B. *UFO*^*Pro-dis-1*^ mutants’ petal count quantified by order of flower emergence. Petal counts are shown in magenta and mean petal counts by flower are depicted by black diamonds.

